# Disruption of fish gut microbiota composition and holobiont’s metabolome by cyanobacterial blooms

**DOI:** 10.1101/2021.09.08.459397

**Authors:** Alison Gallet, Sébastien Halary, Charlotte Duval, Hélène Huet, Sébastien Duperron, Benjamin Marie

## Abstract

**Background:** Cyanobacterial blooms are one of the most common stress encountered by metazoans living in freshwater lentic systems such as lakes and ponds. Blooms reportedly impair fish health, notably through oxygen depletion and production of bioactive compounds including cyanotoxins. However, in the times of the “microbiome revolution”, it is surprising that so little is still known regarding the influence of blooms on fish microbiota. In this study, an experimental approach is used to demonstrate that blooms affect fish microbiome composition and functions, as well as the metabolome of holobionts. To this end, the model teleost *Oryzias latipes* is exposed to simulated *Microcystis aeruginosa* blooms of various intensities in a microcosm setting, and the response of bacterial gut communities is evaluated in terms of composition, metagenome-encoded functions and metabolome profiling.

**Results:** The gut bacterial community of *O. latipes* exhibits marked responses to the presence of *M. aeruginosa* blooms in a dose-dependent manner. Notably, abundant gut-associated Firmicutes almost disappear, while potential opportunists increase. The holobiont’s gut metabolome displays major changes, while functions encoded in the metagenome of bacterial partners are more marginally affected. Bacterial communities tend to return to original composition after the end of the bloom suggesting post-bloom resilience, and remain sensitive in case of a second bloom, reflecting a highly reactive gut community.

**Conclusion:** In the context of increasingly frequent and intense blooms worldwide, results point to the relevance of accounting for short- and long-term microbiome-related effects in fish ecology, with potential outcomes relevant to conservation biology as well as aquaculture.

## Background

Organisms living in lakes and ponds are exposed to cyanobacterial blooms throughout their life (1–4). Blooms are natural events, yet increased eutrophication and global change associated with human activities make them increasingly frequent, abundant and persistent worldwide (5, 6). Cyanobacterial blooms affect the whole ecosystem, including the health of teleost fish which occupy higher trophic levels (1, 2, 7). Cyanobacterial metabolites display a broad range of bioactivities and include various toxins, digestive enzyme inhibitors, antimicrobials, and cytotoxic compounds (8). As a consequence, cyanobacteria cells, extracts and purified cyanotoxins all induce deleterious effects on teleost fishes (2, 9–12). One of the most toxic and frequent cyanotoxins is microcystin-LR (MC-LR), a hepatotoxin that accumulates in fish liver with deleterious consequences for fish physiology and reproductive processes (13, 14). Aquatic animals are exposed to cyanobacteria and their toxins through oral ingestion and transfer absorption through the intestine (13, 15, 16). In this respect, the role of gut-associated microbiota in holobiont’s response is currently underestimated (17). The gut microbiota has recently emerged as a primary target for microbiome-aware ecotoxicological concerns (18–22). However, a limited number of studies have investigated the effect of cyanobacterial blooms on fish gut microbiota (10, 23–25). Whole chemical extracts of few *Microcystis* strains were for example shown to influence the composition of gut bacterial communities of medaka fish in microcosm-based experiments, while MC-LR alone did not (24). This and few other works emphasize that metabolite cocktails and whole cells, rather than toxins alone (microcystins), should be considered for realistic assessment of the microbiome impairs (2, 13, 24), yet the effect of exposure to environmentally relevant levels of cyanobacteria has not been evaluated so far.

In the present study, the impact of a cyanobacterial bloom on the composition and functions of fish gut-associated bacterial communities, and on the metabolite composition in various host tissues is evaluated using an experimental approach. A teleost model fish, the medaka *Oryzias latipes*, has been exposed to three environmentally relevant concentrations of *Microcystis aeruginosa*, the most common bloom-forming cyanobacterium in temperate lentic freshwaters (26). A first 28-days exposure simulated a long bloom event. Because blooms are highly dynamic events in natural systems, post-bloom resilience was investigated. Then, the hypothesis of a priming effect, translating into a lower impact of a second bloom, was tested. To this end, a post-bloom depuration phase was conducted for 4 days, followed by a second exposure to the highest *M. aeruginosa* concentration for 5 days. Bacterial community compositions were characterized using 16S rRNA gene sequencing, and metabolite contents were profiled by LC-MS/MS. Metagenomes of unexposed control fish gut communities were compared to those of specimens exposed to the highest bloom level to compare their respective annotated functions. Bacterial community and metabolites compositions were then compared using a multi-omics approach to identify correlation networks associated with holobiont response. By testing the effect of a cyanobacterial bloom on teleost gut bacterial microbiota, documenting holobiont post-bloom response, and investigating the effect of a second bloom, this study addresses for the first time the dynamics of holobiont response to cyanobacterial blooms.

## Methods

### Experimental design and sampling

Experimental procedures were carried out in accordance with European legislation on animal experimentation (European Union Directive 2010/63/EU) and were approved for ethical contentment by an independent ethical council (CEEA Cuvier n°68) and authorized by the French government under reference number APAFiS#19316-2019032913284201 v1.

Experiments were performed in 10-liter aquaria (microcosms) with 7-months old adult male Japanese medaka fish *Oryzias latipes* provided by the AMAGEN platform (Gif-sur-Yvette, France). Before the whole experiment, five fish were sampled as controls for the histological analyses then fish were pre-acclimatized in clear water (2 weeks) in 15 aquaria, each one containing 8 fish. Prior to the first exposure at day 0, five fish were randomly sampled among the aquaria, and 10 mL of water of each aquarium were pooled, as references of initial fish and water conditions (d0, Fig. 1). Fish were then exposed for 28 days to five treatments: water (control, 0); water containing Z8 medium (27), *i.e*. the medium used to cultivate cyanobacteria (control, Z8); and water containing three environmentally relevant concentrations of live *Microcystis aeruginosa, i.e*. 1, 10 and 100 *µ*g.L^−1^ Chl*a* (1, 10, 100), respectively (d28, Fig. 1). Each treatment was carried out in three aquaria (labelled a, b and c). At d28, four fish, 150 mL of water, bottom-growing biofilms and faeces were sampled in each aquarium. After sampling, remaining fish from each of the three aquaria exposed to one condition were pooled and transferred to a single aquarium, filled with clear water (treatment 0) for a 4-day depuration period of exactly 110 hours (d33, Fig. 1). At d33, three fish and 150 mL of water were sampled in each aquarium. All fish were then exposed in the same aquaria for 5 days (exactly 134 hours) to the highest concentration of *M. aeruginosa*, 100 *µ*g.L^−1^ Chl*a* (d39, Fig. 1). At d39, four fish, 150 mL of water and faeces were sampled in each aquarium. Along the two successive exposures, 10 mL of *M. aeruginosa* culture were sampled every two days but only one sample per week was analysed further. Dataset S1 provides details of sampled individuals and performed analyses.

**Figure 1.**
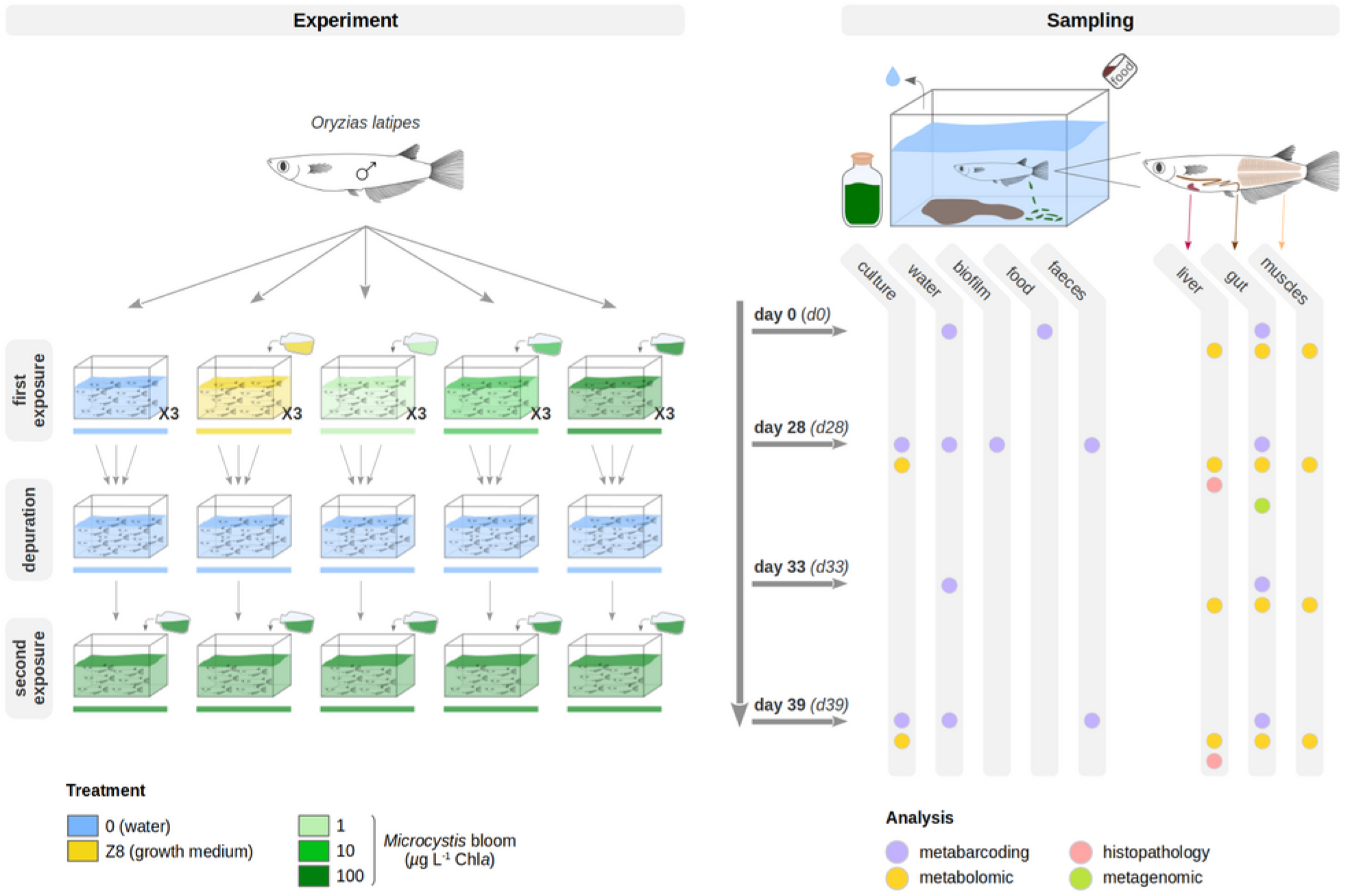
Experimental design and *O. latipes* sampling strategy. After a 14-days acclimatisation period, adult male *O. latipes* were exposed to two successive exposures. During the first 28 days (d0 to d28), fish were exposed to five different treatments, each carried out in three 10-liter aquaria. Treatments consisted of two controls (0, Z8) and three concentrations of *Microcystis aeruginosa* (1, 10 and 100 *µ*g.L^−1^ Chl*a*). After that, a 4-days depuration phase followed (d28 to d33) performed in clear water (0), then fish were exposed again for 5 days (d33 to d39) to *M. aeruginosa* (100 *µ*g.L^−1^ Chl*a*). Colours represents the different treatments: blue = water, yellow = Z8 growth medium, light green = 1 *µ*g.L^−1^ Chl*a*, green = 10 *µ*g.L^−1^ Chl*a*, dark green = 100 *µ*g.L^−1^ Chl*a*. Samples were collected at d0 (day 0), d28 (day 28), d33 (day 33) and d39 (day 39). Further analyses were performed on *M. aeruginosa* culture, water, biofilm, fish food, faeces samples and different tissues (Supplementary Note 1 and Fig. S1a-d). Dataset S2 provides details on sampling counts.

### *Microcystis aeruginosa* production

Blooms were simulated in lab using the non-axenic and easy-to-cultivate *M. aeruginosa* mono-clonal strain PMC 728.11 maintained in the Paris Museum Collection which can produce diverse variants of bioactive metabolites (Supplementary Note 2). The strain was cultivated in Z8 medium at 25 ± 1 °C with a 16h:8h light/dark cycle (at 14 *µ*mol.m^−2^.s^−1^) in 2-liter bottles all along the experiment. The concentrations of *M. aeruginosa* were estimated using Chlorophyll *a* extraction (28) and absorbance measurements as a proxy using a spectrophotometer (Cary 60 UV-Vis, Agilent). Every two days, after water renewal, *M. aeruginosa* was measured in each aquarium using a fluorometer (FluoroProbe III, bbe Moldaenke), and appropriate volume of the culture was added as per the desired final concentrations (1, 10 or 100 *µ*g.L^−1^ Chl*a*).

### Monitoring of experimental parameters

Every second day throughout the experiment, water parameters were monitored (pH, temperature, conductivity, nitrates and nitrites), aquaria were cleaned (faeces removed by aspiration), half of the water was replaced with freshwater. On the same timeline, *M. aeruginosa* concentrations were measured and adjusted to maintain bloom levels in exposed treatments, and 1 milliliter of sterile Z8 medium was added in the Z8 treatment. Fish were exposed to constant temperature (23 ± 1 °C), pH (7.5 ± 0.1) and conductivity (234 ± 22 *µ*S.cm^−1^), to low levels of nitrates (≤ 1 mg.L^−1^) and nitrites (≤ 4 mg.L^−1^), to a 12h:12h light/dark cycle, and were fed twice daily with Nutra HP 0.3 (Skretting, Norway). Microcystin (MC) concentration was monitored on a regular basis in *M. aeruginosa*-containing treatments and quantified using enzyme-linked immunosorbent assay (ELISA) analyses (Microcystins-ADDA SAES ELISA, Eurofins Abraxis). Each sample was analysed in duplicates and MC concentrations were determined according to the MC-LR response standard curve.

### Fish, *M. aeruginosa* culture, water, biofilm, food and faeces processing

Fish were anesthetized in 0.1% tricaine methanesulfonate (MS-222; Sigma, St. Louis, MO) buffered with 0.1% NaHCO_3_ and sacrificed. Whole guts, muscles and livers were dissected, flash-frozen in liquid nitrogen and stored at -80 °C. For histopathological examinations, livers from fish sampled before the whole experiment and at d28 and d39 (one fish per aquarium) were dissected, fixed in Davidson fixative as previously described(13), maintained for 24h at 4 °C then dehydrated in 70% ethanol and conserved at 4 °C. Livers samples were then embedded in paraffin and blocks were cut into 4 *µ*m thick sections, stained with Hematoxylin-Eosin-Saffron (HES), Periodic Acid Schiff (PAS) and Perls Prussian blue, and observed under photonic microscope (Zeiss, Germany). Aquarium water samples were filtered on a 0.22-*µ*m filter (Nucleopore Track-Etch Membrane) and frozen. *M. aeruginosa* culture (4 mL) and biofilm samples were centrifugated (10 min, 10 °C, 3,220 g) and pellets were frozen. Faeces pellets were directly frozen. A food sample was kept for DNA extraction.

### Metabolites extraction

Metabolite contents were extracted from fish livers, guts and muscles, *M. aeruginosa* cultures, and biofilms. Prior to sonication, muscles were freeze-dried then ground using a bead beater (TissueLyser II, Qiagen) while cultures and biofilms were only freeze-dried. Samples were weighted, then sonicated in 75% methanol (1 mL per 100 mg of tissue, 3x, on ice) and centrifuged (10 min, 4 °C, 15,300 g). Supernatants containing metabolite extracts were kept at -20 °C for mass spectrometry analyses. All pellets were discarded, except gut pellets dried and kept at -80 °C to perform a subsequent DNA extraction on the same gut tissue.

### Mass spectrometry data processing and analysis

Each metabolite extract from fish livers, guts and muscles was analysed by Ultra high-performance liquid chromatography (UHPLC; ELUTE, Bruker) coupled with a high-resolution mass spectrometer (ESI-Qq-TOF Compact, Bruker) at 2 Hz speed, on simple MS mode then on broad-band Collision Ion Dissociation (bbCID) or autoMS/MS mode on the 50-1500 *m*/*z* range. Three feature peak lists were generated from MS spectra within a retention time window of 1-15 minutes and a filtering of 5000 counts using MetaboScape 4.0 software (Bruker). The three peak lists consisted of the area-under-the-peaks of extracted analytes from the three tissues (gut, liver, muscle) sampled at d28, d33 and d39, resulting in 1672, 909 and 3127 analytes, respectively. The genuine metabolite content of the culture was investigated on metabolite extracts using LC-MS/MS approach, combined with molecular network analysis and metabolite annotation using a cyanobacterial metabolite reference database, as previously described (29). Prior to analyses, Pareto scaling was applied on the datasets. Principal Component Analyses (PCA) were performed to compare the metabolite composition among groups using the *mixOmics* package (30) in R 4.1.0 (R Core Team, 2021). The variance among groups was compared conducting PERMANOVA (999 permutations) based on euclidean distance with the *vegan* (31) followed by Bonferroni-adjusted pairwise comparisons with the *RVAideMemoire* (32).

### DNA extraction

DNA was extracted from gut, culture, biofilm and faeces pellets, water filters and food using the ZymoBIOMICS DNA Miniprep kit (Zymo Research, California). Prior to DNA extraction, all pellets were re-suspended in Eppendorf tubes with 750 *µ*L of the ZymoBIOMICS™ lysis solution, then the contents were transferred to the ZR BashingBead™ lysis tubes. Water filters were cut into pieces then transferred to the ZR BashingBead™ lysis tubes. All steps were conducted following the manufacturer’s instructions except for mechanical lysis, achieved on a bead beater (TissueLyser II, Qiagen) during 6×1 min. An extraction blank was performed as a control. The quality and quantity of the extracted DNA was tested on Q-bit (Thermo).

### Bacterial 16S rRNA gene sequencing and analyses

The V4-V5 variable region of the 16S rRNA gene was amplified using 479F (5’-CAGCMGCYGCNGTAANAC-3’) and 888R primers (5’-CCGYCAATTCMTTTRAGT-3’) (33), and sequenced (Illumina MiSeq paired-end, 2×250 bp, GenoScreen, France). Paired-end reads were demultiplexed, quality controlled, trimmed and assembled with FLASH (34). Sequence analysis was performed using the QIIME 2 2020.11 pipeline (35). Chimeras were removed and sequences were trimmed to 367 pb then denoised using the *DADA2* plugin, resulting in Amplicon Sequence Variants (ASVs) (36). ASVs were affiliated from the SILVA database release 138 (37) using the *feature-classifier* plugin and *classify-sklearn* module (38, 39). Sequences assigned as Eukaryota, Archaea, Mitochondria, Chloroplast and Unassigned were removed from the dataset then the sample dataset was rarefied to a list of 6,978 sequences. Alpha- and beta-diversity analyses were performed using the *phyloseq* (40), *vegan* and *RVAideMemoire* packages in R. Linear mixed models (LMMs) were used to compare species richness among the five treatments and the three replicate aquaria within each treatment at d28, using the *MuMIn* (41) and *lmerTest* (42) R packages. We adapted the LMMs to the non-independency of individuals within each replicate, and defined the replicates as random effects and the five treatments as fixed effects, according to the formula Y ∼ treatment + (1 | treatment : aquarium). Principal coordinates analyses (PCoA) based on weighted and unweighted UniFrac distances were performed to examine the dissimilarity of bacterial composition between groups. Among- and within-group variance levels were compared using PERMANOVA (999 permutations) and PERMDISP (999 permutations), respectively. Differentially abundant taxa across groups were identified using the linear discriminant analysis (LDA) effect size (LEfSe) tool (43) in the Galaxy workspace (44) (http://huttenhower.sph.harvard.edu/lefse/). Default parameters were applied except for the LDA score threshold and for the different multi-class strategy (one-against-all).

### Microbiome-metabolome integrative analysis

The integration of datasets, *i.e*. the area-under-the-peaks in metabolite profiles and the ASV counts describing the bacterial communities in the same sample, was performed using the *mixOmics* package in R. Pareto scaling was applied on the metabolome data, and a centred log-ratio transformation then a prefiltering keeping only abundant ASVs, (*i.e*. representing at least 1% of the reads in at least one sample), were applied on the microbiome data. Following unsupervised analyses on each dataset, completed to explore and visualize any similar changes according to treatments, the integration was carried out. A supervised Projection to Latent Structures Discriminant Analysis (PLS-DA) was performed using DIABLO (*Data Integration Analysis for Biomarker discovery using Latent cOmponent*) (45), enabling to identify highly-correlated variables (metabolites and ASVs) also discriminating the different treatments. The integration of both datasets was realised using the full weighted design matrice and the *block.plsda* function implemented in *mixOmics*. The *plotDiablo* function enabled to check the well maximized covariation between datasets by displaying a Pearson correlation score. Then, relevance networks displaying the most discriminant covariates (metabolites, ASVs) were produced using the *network* function with Pearson correlation cut-offs (46).

### Metagenomic sequencing and analysis

Shotgun metagenome sequencing was performed on DNA from 10 gut samples collected at d28, 5 in treatment d28_0 and 5 from d28_100 (Illumina HiSeq, 2×150bp, GenoScreen, France). Reads corresponding to animal sequences were identified by aligning each dataset against *Oryzias latipes* available at the NCBI, using BBMap/bbsplit (47), and discarded. Remaining reads from each sample were assembled using metaSPAdes with default parameters (48). Scaffolds were first taxonomically annotated using Contig Annotation Tool (CAT) (49) and Kaiju (50) allowing to detect sequences from *O. latipes* retroviruses and *Microcystis* genome which were then discarded. All scaffolds were clustered using MyCC (51) (k-mer size = 4, minimal sequence size = 1000) and bins were taxonomically annotated using Bin Annotation Tool(49). Completeness of bins was assessed using CheckM (52). Relative abundance of bins in each sample were also determined using BBMap. Significant bins between the two treatments were determined using Wilcoxon rank-sum test. Finally, coding sequences predicted by Prodigal (53) were functionally annotated using eggNOG-emapper (54). Resulting KEGG annotations were used as input to MinPath (55) in order to obtain the complementary MetaCyc pathway information.

## Results

### Monitoring of experiments

*Oryzias latipes* fish were maintained in suitable and stable conditions, and no fish died during the whole experimentation (Fig. 1 and Dataset S2). During the first 28-days long exposure (d0-d28), concentrations of *Microcystis aeruginosa* in the three treatments (d28_1, d28_10 and d28_100) corresponded to expected levels, 1.0 ± 0.2, 10.0 ± 0.8 and 100.2 ± 7.8 *µ*g.L^−1^ Chl*a*, respectively. Microcystin levels in water were 0.4 ± 0.0 and 10.4 ± 2.1 *µ*g MC-LR eq.L^−1^ in the d28_10 and d28_100 treatments, respectively, while microcystin was below detection level (< 0.15 *µ*g.L^−1^) in d28_1. During the second *M. aeruginosa* exposure (d33-d39), fish were exposed to 102.8 ± 2.9 *µ*g.L^−1^ Chl*a* and 11.2 ± 2.8 *µ*g MC-LR eq.L^−1^. The metabolic content of *Microcystis aeruginosa* cultures was examined (see Supplementary Note 2 and Table S1). Histopathological analyses did not reveal noticeable visual differences in fish liver tissue, with little to no carbohydrate reserves, and no lipofuscin and macrophagic hemosiderin.

### Diversity and composition of the gut bacterial microbiota after 28 days of exposure

At d28, gut communities display between 42 and 219 ASVs with higher average bacterial richness (136 ± 56 ASVs) and evenness (0.556) reported in fish guts exposed to d28_Z8, and lower average bacterial richness (82 ± 25 ASVs) and evenness (0.391) in the treatment d28_0 (Dataset S4). Gut-associated species richness was highest in the d28_Z8 treatment (LMM, *p* < 0.05), while no differences are observed among replicates within each treatment (LMM, *p* > 0.05). The Shannon index increases slightly with *Microcystis* concentration (d28_1: 2.09, d28_10: 2.14, d28_100: 2.28). Visual comparisons on individual plots of principal coordinates analyses (PCoA) based on the weighted or unweighted UniFrac distances suggest changes occur in terms of both abundances as well as community membership when fish are exposed to *M. aeruginosa* (d28_1, d28_10, d28_100) or d28_Z8 compared to d28_0 (Fig. 2a,b). Bacterial community composition appears different among treatments (PERMANOVA, weighted UniFrac, *p* < 0.001), notably between d28_100 and the other four treatments (*p* < 0.01), as well as between d28_0 and d28_Z8, d28_1 and d28_100 (*p* < 0.02), but not between d28_Z8, d28_1 and d28_10 (*p* > 0.23). In addition, levels of variance in the different treatments are not significantly different (PERMDISP, *p* > 0.19).

**Figure 2.**
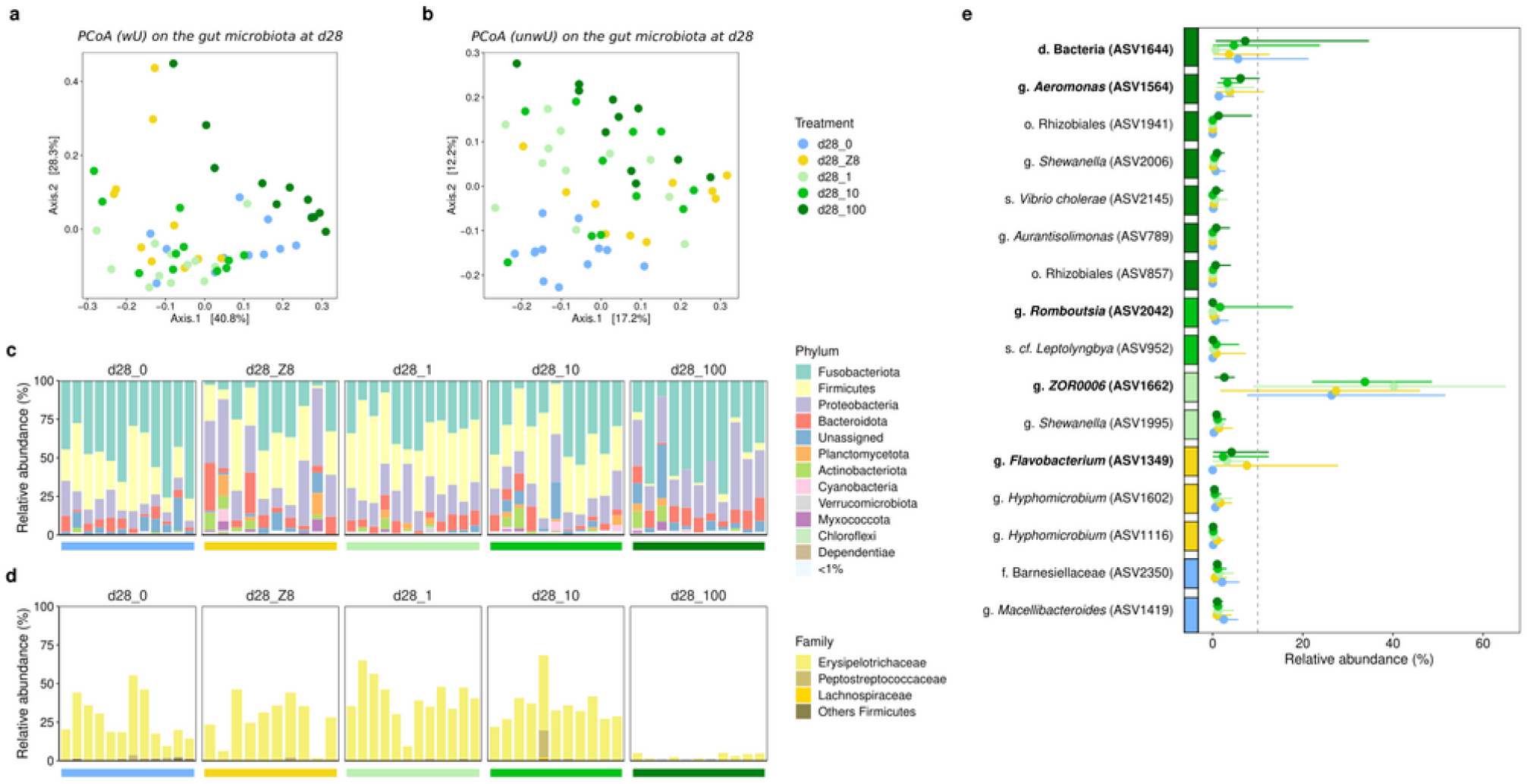
Changes in the composition of gut bacterial communities after 28 days of exposure. (a-b) PCoA using the weighted (a) or unweighted (b) UniFrac distances on fish gut bacterial community in the five treatments. (c) Relative abundance of bacterial phyla across treatments. (d) Relative abundance of Firmicutes members (family level). Firmicutes are mainly represented by the Erysipelotrichaceae family, and a single ASV (ASV1662). Coloured horizontal bars represent the different treatments (see Fig. 1). (e) Significant ASVs from the Linear discriminant analysis (LDA) effect size (LEfSe) with a LDA score above 3.5, and their relative abundances across d28 treatments. The coloured boxes on the y-axis represent the treatment where each ASV is most abundant; dots represent the average relative abundance; line spreads over the range of observed values. Only dominant ASVs, *i.e*. representing at least 10% of reads in at least one sample were further considered.

In treatment d28_0 (water-exposed control), Fusobacteriota (50 ± 16%), Firmicutes (28 ± 15%), Proteobacteria (10 ± 6%) and Bacteroidota (5 ± 3%) dominate the microbiota, altogether representing 92.8 ± 6.9% of the reads. Relative abundances of these phyla are different among other treatments. Notably, Firmicutes are far less abundant in d28_100 (3 ± 2%) compared to other treatments (d28_0: 28 ± 15%, d28_Z8: 28 ± 15%, d28_1: 41 ± 14%, d28_10: 36 ± 12%) (Fig. 2c).

Many ASVs display significant differences across treatments in their relative abundances (Fig. 2e and Fig. S2a). We focus on the six dominant ASVs, *i.e*. representing at least 10% of reads in at least one sample, accounting from 11 to 72% of the reads. ASV1349, ASV1564 and ASV2363 exhibit lower relative abundances in d28_0 (average below 0.7%). ASV1349 (*Flavobacterium*) and ASV1564 (*Aeromonas*) are more abundant in d28_Z8 (7.6 ± 10% and 3.8 ± 3.5%, respectively) and d28_100 (4.2 ± 4.1% and 6.2 ± 2.9%, respectively). ASV2363 (*Reyranella*) is also more abundant in d28_100 (5.8 ± 11.1%). ASV1662 affiliated to the genus *ZOR0006*, is the main Firmicutes (96-100% of all Firmicutes reads), and as previously mentioned, is least abundant in d28_100 compared to other treatments (Fig. 2d). ASV2042 (*Romboutsia*), dominant in a single sample (17.8%) while below 3.5% in all others, was not further considered. Finally, ASV1644 (unassigned Bacteria) displays similar abundances in treatments d28_0 (5.6 ± 6.0%) and d28_100 (7.2 ± 9.7%) but is much less abundant in treatment d28_1 (0.7 ± 1.2%). Aside from the dominant ASVs that vary, ASV1620 (*Cetobacterium*) displays non-significant differences in abundance among treatments, representing 48 ± 16% of reads in d28_0 and average 27% to 46% of reads in other treatments.

### Metagenome-based comparison of gut communities in d28_0 and d28_100

Metagenome datasets obtained from 5 fish guts from d28_0 and 5 others from d28_100 yielded negligible amounts of Archaea and non-fish Eukaryota sequences. Among the 32 bacterial bins obtained, 11 are found to be differentially abundant between the two treatments (*p* < 0.05, Wilcoxon rank-sum test) (Fig. S2b). Eight are more abundant in d28_100, including seven assigned to the same taxa as ASVs (*Flavobacterium, Aeromonas, Gemmobacter*, Rhizobiales), and vary in the same way as ASV counts (Fig. S2a,b). Of the 3 bins found significantly more abundant in d28_0, bin24 is affiliated to the Firmicutes, with a majority of coding sequences associated to *ZOR0006* (58.06%), the genus corresponding to aforementioned ASV1662. Gene functions were investigated by analysing enzymes (KO) potentially produced by bins and their associated pathways. Bins can be distinguished into three groups according to Wilcoxon rank-sum test performed (*p*-value fixed at 5%) between treatments d28_0 and d28_100 (Fig. S2c). Only 0.3% of enzymes are specific to the three bins that are more abundant in d28_0. Altogether, bin24 (*ZOR0006*), bin22 (Bacteroidales) and bin11 (Porphyromonadaceae) count 6 specific enzymes, including one involved in lactose degradation (KO2788). On the other hand, 4.3% of enzymes are specifically found in the 8 bins that are enriched in d28_100 (namely bin9, bin10, bin12, bin16, bin19, bin23, bin25, bin26). These represent 95 specific enzymes, including 14 involved in the biosynthesis of secondary metabolites, 10 in porphyrin and chlorophyll metabolism, and 6 in the biosynthesis of cofactors. The analysis of clusters of orthologous groups (COGs), regardless of their annotation status, reveals that 963 of them (4.4% of the total COGs) are specific to the three bins that are the most abundant in d28_0, and that 2316 others (10.7% of the total gene families) are specific to the 8 bins that are the most abundant in d28_100. However, one should notice that most of these COGs carry no known functional annotation, compromising our understanding of the influence of gene content variations on microbiota functioning based on metagenomic assumption.

### Gut metabolome variation and integration with gut microbiota composition after 28 days of exposure

A total of 1,674 metabolites were detected across gut samples at d28. The PCA analysis separates the metabolite profiles of fish exposed to the different *M. aeruginosa* levels (d28_1, d28_10, d28_100) along the first axis, while the second axis mostly separates d28_0 from other treatments (Fig. 3a). The gut metabolite composition is different among treatments (PERMANOVA, euclidean distance, *p* < 0.001), between d28_0 and the three *M. aeruginosa* treatments (*p* = 0.01), and between d28_100 and all treatments (*p* < 0.03) but d28_10 (*p* = 0.19). Treatments d28_Z8, d28_1 and d28_10 do not display differences (*p* > 0.3). No significant variation is observed among treatments in livers (*p* = 0.13) (Fig. S3a). In muscles, differences occur (*p* < 0.001) especially between d28_0 and both d28_Z8 and d28_1 (*p* = 0.01), and between d28_100 and all other treatments (*p* = 0.01) except d28_0 (*p* = 0.33) (Fig. S3b).

**Figure 3.**
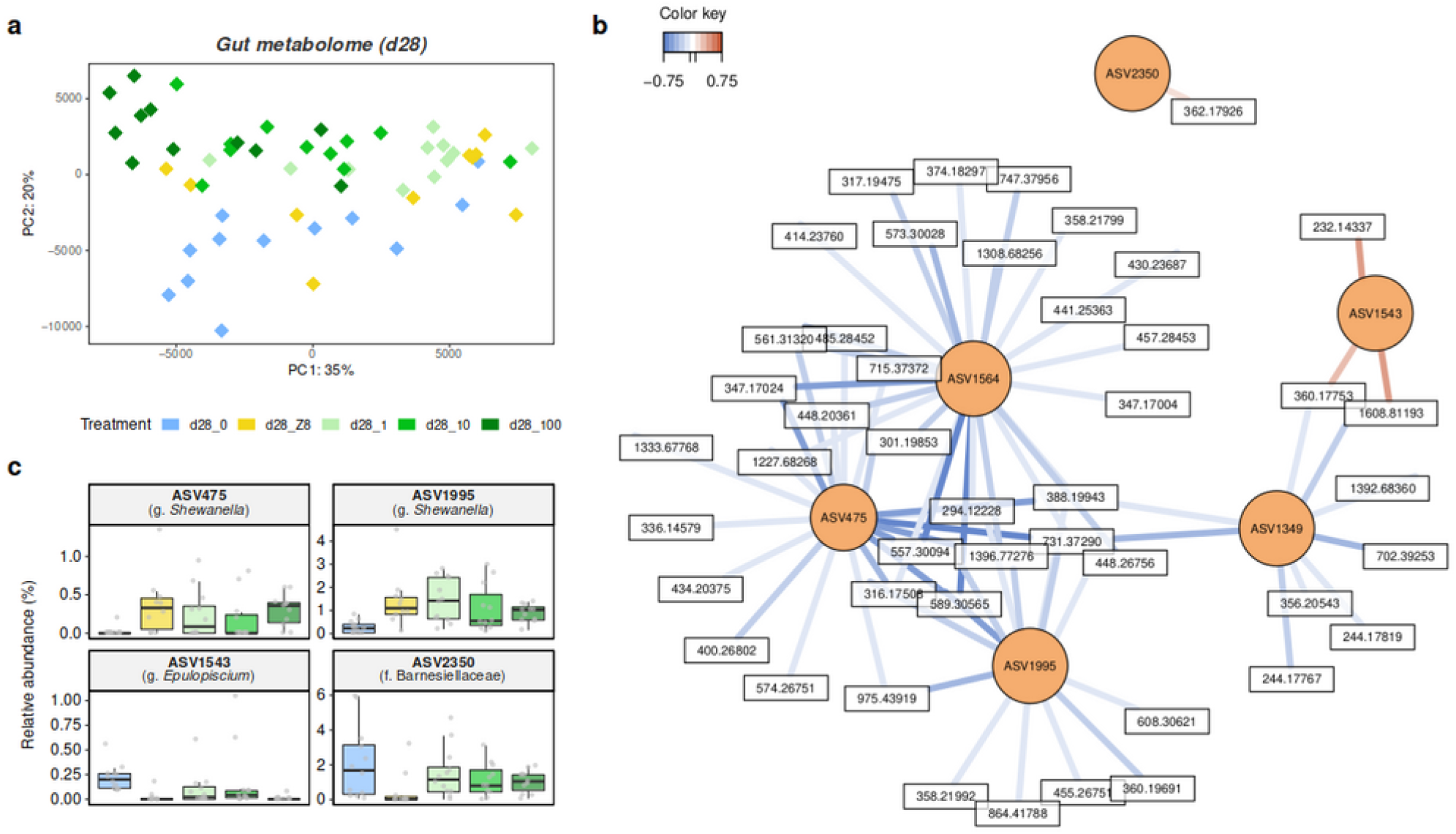
Modification of the gut metabolome composition associated with bacteria changes. (a) Principal component analysis (PCA) representing gut metabolite profiles at d28. (b) Relevance network analysis illustrating the most correlated metabolites (in white) and ASVs (in orange) discriminating among the five treatments. Only variables (ASVs, metabolites) with correlation values above ±0.695 are displayed. Pearson correlations between covariate metabolites and ASVs are represented by coloured segments (blue: negative association, red: positive association). (c) Relative abundances of the four discriminant ASVs, displayed in the relevance network and not mentioned in previous analyses.

A joint analysis of gut metabolites and bacterial communities was performed, as described by Singh and colleagues (45). The application of this multi-omics method has required the initial development of a new protocol for the extraction of both metabolites and DNA from the single gut samples of very small organisms. The resulting combined dataset consists of two matrices of the 70 abundant ASVs, *i.e*. representing at least 1% of reads in at least one sample, and the 1674 metabolites, sufficiently well correlated together (76%) to explore associations between ASVs and metabolites. The relevance network illustrates correlations between the most highly associated ASVs and metabolites discriminating among the five treatments. Four ASVs (ASV1349, ASV1564, ASV475, ASV1995) appear negatively correlated with several metabolites and are the least abundant in d28_0 (Fig. 2e and Fig. 3b,c). Differently, the two other ASVs, namely ASV1543 (*Epulopiscium*) and ASV2350 (Barnesiellaceae), are positively correlated with few metabolites and are the most abundant in d28_0 (Fig. 3b,c). Unfortunately, only few of the metabolites presenting high correlation with abundant ASVs could be annotated, and these mainly corresponded to amino acids or small peptides (Dataset S5).

### Diversity and composition of the gut bacterial microbiome and metabolome after depuration (d33) and a second exposure (d39)

After d28, fish from all 5 treatments were transferred to clean water (Fig. 1). At d33, some specimens were sampled, while others were transferred to a second exposure to *M. aeruginosa* (100 *µ*g.L^−1^ Chl*a*), then sampled at d39. The species richness of gut-associated communities decreases between d28 and d33 (over 82 versus 50 ± 24 ASVs), then increases at d39 (76 ± 22 ASV). The PCoA discriminates between gut communities from fish exposed to 100 *µ*g.L^−1^ Chl*a* (d28_100 and d39) and other treatments (Fig. S4a). Community compositions differ among treatments (PERMANOVA, weighted UniFrac, *p* < 0.001). Treatment d33 differs from all other treatments (*p* < 0.05) except for d28_0 (*p* = 0.063); in other words, bacterial communities at d33 are mostly similar to those observed in the d28_0 treatment. Communities at d39 differ from all other treatments (*p* < 0.05), except for d28_100 (*p* = 0.063); indicating that bacterial communities at d39 are similar to those from the d28 treatment exposed to the highest bloom intensity. Variance differs among treatments (PERMDISP, *p* < 0.01), particularly between d33 and all other treatments (*p* < 0.05), and between d39 and treatments d33 and d28_Z8 (*p* < 0.05), with d33 and d39 displaying lower variance. Some phyla are present in similar abundances at d33 and d39, including Fusobacteriota (62 ± 8% and 66 ± 14%, respectively), Proteobacteria (12 ± 6% and 19 ± 9%) and Bacteroidota (9 ± 3% and 7 ± 3%, Fig. 4a,b). Firmicutes, again mostly consisting of ASV1662, are present at d33 and almost absent at d39 (15.4 ± 6.4% versus 0.8 ± 1.8%). Other significant changes have been observed in specific ASVs between d33 and d39 (Fig. S5a) among which seven were already observed to vary between the different d28 treatments (Fig. 2e and Fig. S2a). Interestingly, ASV1620 (*Cetobacterium*), that was found abundant and stable among the different d28 treatments, is still remarkably stable at d33 and d39 (58.8 ± 7.4% and 61.7 ± 13.2%).

**Figure 4.**
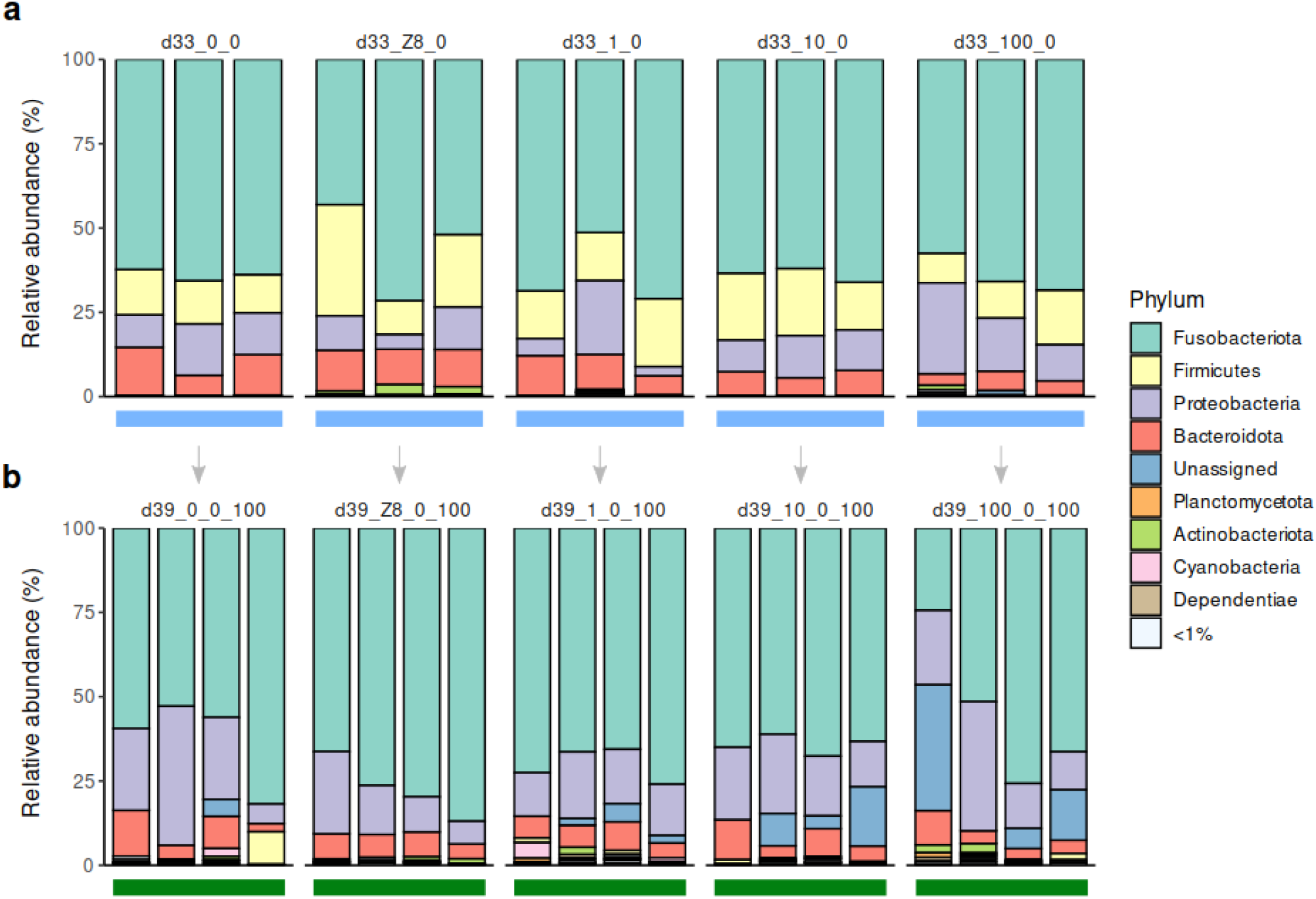
Depuration and a second *Microcystis* bloom impact the gut microbiota community. Relative abundance of gut bacterial phyla at d33, after 4 days in water following the d28 treatment (a), and at d39 after 5 additional days exposed to *M. aeruginosa* (b). Coloured horizontal bars represent the different imposed treatments: blue = water, dark green = 100 *µ*g.L^−1^ Chl*a*.

The gut metabolite profiles are significantly different among treatments (PERMANOVA, euclidean distance, *p* < 0.001, Fig. S4b), especially between d33 and all treatments (*p* < 0.05) except d28_Z8 and d28_1, as well as between d39 and all others (*p* < 0.05) except d28_Z8 and d28_10. Contrary to bacterial microbiota composition, the d33 metabolomes are overall different from d28_0, and those of d39 are different from d28_100. A good correlation (81%) was observed between the gut ASVs and the metabolites discriminating between d33 and d39 when investigating the two matrices, containing 31 abundant ASVs and 1674 metabolites, respectively. The relevance network notably revealed two ASVs, belonging to genus *Vibrio* (ASV1406, ASV2161), that are negatively correlated with numerous metabolites, while two ASVs, belonging to genus *Reyranella* (ASV1030, ASV2363) appear positively correlated with numerous metabolites (Fig. S5b). No differences occur among treatments in liver samples (*p* > 0.05, Fig. S4c). In muscles, metabolomes at d33, d39 and d28_100 appear similar (p > 0.10), but different from all other treatment groups (Fig. S4d).

### Occurrence of dominant ASVs in compartments other than gut

Dominant ASVs were searched for in fish food, *M. aeruginosa* culture, water, biofilm, and faeces (Fig. S6). ASV1620 (*Cetobacterium*) is most abundant in guts (44.3 ± 20.4%) then in faeces (28.6 ± 15.3%), congruent with its best matches in the database which are fish gastrointestinal bacteria (zebrafish, Nile Tilapia). ASV1349 (*Flavobacterium*) and ASV1564 (*Aeromonas*) are most abundant in faeces (8.8 ± 9.6% and 7.8 ± 2.8%, respectively), then guts (2.1 ± 4.4% and 3.4 ± 2.6%, respectively) and biofilms (2.0 ± 3.3% and 3.4 ± 2.5%, respectively). This is congruent with their respective BLAST_N_ hits with bacteria from fish intestinal tracts or aquatic environments. ASV1644 (unassigned Bacteria) is also most abundant in faeces (10.7 ± 7.4%), then in guts (3.7 ± 6.6%), but could not be assigned taxonomically. The Firmicutes ASV1662 (*ZOR0006*) mainly occurs in guts (19.3 ± 16.9%) and is slightly abundant in faeces (1.2 ± 1.4%), in accordance with the habitat of its closest matches, namely fish intestinal bacteria. Finally, ASV2363 (*Reyranella*) is most abundant in biofilms (4.4 ± 8.7%) compared to other compartments (water: 2.2 ± 6.3%; faeces: 1.3 ± 2.7%; guts: 1.2 ± 4.1%), and is related to various environmental bacteria.

## Discussion

Results from the 28-days exposure indicate that *Microcystis aeruginosa* blooms modify the fish gut bacterial microbiota compositions and have different effects on different taxa. Firmicutes, a phylum commonly found in gut bacterial communities of vertebrates and very likely implied in host metabolism processes (56–58), appear particularly sensitive as they decrease sharply upon exposure to the highest bloom intensity. Firmicutes were largely represented by a single bacterium (Erysipelotrichaceae_*ZOR0006*) that was almost absent outside of gut samples, suggesting being an indigenous and resident symbiont of *O. latipes* gut. Kaakoush and colleagues (59) have previously discussed the central role of Erysipelotrichaceae in host lipid metabolism and health in relation to diet specificities. The *ZOR0006* genus has already been observed to decrease in fish gut microbiota after exposing zebrafish 21 days to high concentrations of antibiotics or fungicides (60, 61). Interestingly, the drop of *ZOR0006* abundance observed in this study occurs at a level between 10 and 100 *µ*g.L^−1^ Chl*a*. Together with the observation of the greater influence of the highest bloom condition on the whole gut community, this suggests the existence of a cyanobacterial bloom threshold above which the gut microbiota composition is particularly altered. On the other hand, some abundant gut bacteria appear stable throughout the bloom, and even during the depuration and second exposure, the most remarkable being the bacterium affiliated to *Cetobacterium* (Fusobacteria). This genus is generally associated with healthy fish microbiota and notably contributes to host health as a B_12_ vitamin producer (62–64). *Cetobacterium* is most likely a fish gut resident and has previously been reported stable in medaka fish upon exposure to pure microcystin-LR and cell extract of the *Microcystis* strain (24), supporting its maintenance through the exposure to cyanobacterial bloom and respective metabolites. Finally, relative abundances of other bacteria increase during the *Microcystis* bloom. These may be transient gut bacteria that might originate from grazed biofilms and proliferate once established in the gut, or be rare gut taxa that can take advantage of peculiar conditions to proliferate. In our study, some of these opportunistic bacteria, corresponding to *Flavobacterium, Aeromonas* and *Shewanella*, are common inhabitants of fish guts or the environment, or potential fish gut pathogens according to the literature (65–67). A similar increase of opportunistic bacteria was recently documented in guts of zebrafish exposed 96h to *M. aeruginosa* (10). As previously shown for several metabolite mixtures from cyanobacterial cell extracts (24), exposure to whole cyanobacterial cells thus has a major impact on gut community compositions, indicating that fish microbiota might be impacted during the bloom as well as after bloom senescence which causes the release of the cyanobacterial cell contents into the surrounding water (68). However, only the highest concentration of the bloom and/or its cell extract, that was explored in the present study, could induce most evident microbiota changes, suggesting that the microbiota responsiveness might be dose dependent. The cyanobacterial strain PMC 728.11 contains various secondary metabolites that may be responsible for variations in fish gut microbiota. Among those already identified and specifically those potentially produced by the strain (24, 69), cyanopeptides, such as aerucyclamides and bacteriocins, are thought to exhibit potent antimicrobial or cytotoxic bioactivities (70, 71), and could directly impact the microbiota during *Microcystis* cell digestion into the intestine lumen.

Metagenome-based investigation confirms the variations of taxa abundances, but very few taxa-specific known (*i.e*. annotated) functions were identified for the bacteria that displayed the highest abundance variations, including for the Firmicutes. This is at first consistent with the common claim that a change in microbiota composition does not necessarily imply a change in the functions as estimated by gene content (72, 73). However, similar genes are not necessarily expressed in the same way in different bacterial taxa, and thus phenotypes might still change dramatically despite the potential occurrence of similar gene contents. Indeed, variations in metabolite profiles reveal obvious functional variations induced by the different treatments, in particular changes in gut metabolite composition associated with increasing bloom concentrations. These changes could thus be related to unannotated COGs, much more numerous among the differentially abundant bacteria compared to annotated KO, or could result from variations in the expression of metabolic pathways in the holobiont. Whatsoever, the remarkable correlation observed between some dominant bacteria and many metabolites supports that the response of gut microbiota composition and that of the holobiont’s gut metabolome are linked. Gut community disruption is associated with metabolic changes, especially in the presence of higher bloom levels, suggesting the occurrence of a potential dysbiotic state. However, this crosstalk between gut bacteria and metabolites remains difficult to characterize. For example, the presence of pathogens enhanced by bloom-induced toxicological impairs of fish physiology could secondarily affect the holobiont gut metabolism. Alternatively, the variation of the quantity of some gut metabolites could allow the development of opportunistic pathogens, as reported in mammals (74, 75). Interestingly, variations of metabolite composition in muscles and livers remains limited in comparison to those observed in the gut, implying that the gut metabolome compartment is more responsive to the *Microcystis aeruginosa* exposure. This observation is congruent with the fact that the gut is more directly exposed to the surrounding environment, through ingestion of water that contains various organisms and their associated compounds, and emphasizes the relevance of investigating the gut microbiota, as the gut epithelium sits at the interface between the host and its environment (21, 22). Changes in microbiome and overall holobiont functions seem to be deeply linked, despite that the respective contribution of the host and the bacteria community cannot yet be disentangled and will require further dedicated functional analysis (76).

After the first 28-days bloom, depuration in clear water seems to have restored, to a certain extent, gut microbiota compositions. Indeed, compositions at d33 tended to resemble those of naïve specimens, not previously exposed to *M. aeruginosa* or to Z8 (d28_0). This indicates that the gut community composition is resilient, even over a relatively short time. The following second high-intensity bloom on the other hand yielded bacterial communities resembling those of specimens exposed to the first high-intensity bloom (d28_100), including the disappearance of Firmicutes and the stability of *Cetobacterium*. This indicates that a short duration bloom (5 days) already has a strong effect. Gut communities thus quickly respond to the presence or absence of *M. aeruginosa* and associated bioactive compounds. Effects of the first and the second bloom are however not strictly identical. Firmicutes for example show reproducible behaviour, disappearing upon first and second bloom, and seem to quickly re-establish post-bloom, suggesting they are not resistant, but remarkably resilient. However, the relative abundance of other taxa (including *Flavobacterium, Reyranella, Shewanella*), whose abundance increased upon the 28 days bloom, did not increase during the second 5-days exposure. It remains here difficult to compare variations of most non-dominant bacteria between d33 and d39 because of the substantial inter-individual variation of microbiota composition, combined with the limited sample size in terms of individual number per condition, compared to d28 conditions. Stress associated with specimen handling, the transfer of fish to newly cleaned aquaria after 28 days, has previously been referenced as a moderate, but possible, source of stress (77) and also could be involved to a certain extent in some of the differences. Contrary to gut community compositions, the metabolite composition in fish guts after the depuration and the second exposure did not tend towards those observed in unexposed and highest bloom at d28, respectively. So, metabolite compositions do not linearly follow the trends found in community compositions. This also suggests that the depuration and the second exposure may have been too prompt to induce major shifts in the gut metabolome, as these compartments may present different kinetics. Thus, the dynamics of gut microbiota and metabolome are very likely not identical, and the microbiota composition may somehow respond to changes faster than the metabolome. If confirmed by further investigation, this would suggest that very short blooms occurring in nature, despite their temporary spectacular effect on microbiota composition, may have more limited functional consequences for the holobiont homeostasis (78, 79).

## Conclusions

Overall, this study emphasizes that cyanobacterial blooms should be considered as potent dysbiosis-triggering events for fish gut microbiota and holobiont functions. Additionally, the drop of Firmicutes abundance induced by *M. aeruginosa* bloom which threshold level would be comprised between 10 and 100 *µ*g.L^−1^ Chl*a*, together with the observation of a greater influence of higher bloom condition (100) on the overall gut community, support the existence of a notable tipping point for the responsiveness of gut microbiome to cyanobacterial bloom intensity. This finding could have important eco(toxico)logical consequences, as these levels are commonly reached in natural ecosystems during typical *Microcystis* bloom episodes worldwide, suggesting that destabilization of fish gut communities might be a very common event (26). In nature, cyanobacterial blooms are nowadays becoming increasingly frequent (3), and sometimes persistent among seasons. Freshwater fish, especially those living in shallow water ponds, face numerous bloom episodes of varying durations during their lifetime. Consequences of iterative (up to chronic) exposures on the holobiont should be explored. Indeed, successive blooms without enough recovery time could induce a cumulative drift of the gut microbiota and metabolome, leading to suboptimal states that may lead to host health impairment (80). This phenomenon could be of major significance for fish health in eutrophic natural ecosystems as well as aquaculture ponds where cyanobacteria often proliferate. Testing the effects of a single live *M. aeruginosa* strain administrated by simple balneation on a model teleost fish is a first step towards understanding bloom effects on fish microbiota and holobiont health. In the future, particular attention should also be paid to the natural diversity of cyanobacterial within blooms, as successive blooms often involve different species that produce different metabolite cocktails (81).

## Declarations

### Ethics approval and consent to participate

Experiments were conducted according to best practices within the framework of an authorized program (APAFiS#19316-2019032913284201 v1) approved by Ethical Committee of the National Museum of Natural History (Paris) CEEA n°68.

### Consent for publication

Not applicable.

### Availability of data and materials

16S rRNA gene sequencing and shotgun raw data are deposited into the GenBank SRA database under the BioProject PRJNA746242 (samples SAMN20080275 to SAMN20080438). Files with metabolite tables from mass spectrometry analyses, and the rarefied table of ASVs will be available upon acceptance of the manuscript. The detailed protocol for the extraction of metabolites and DNA will be deposited on protocols.io upon acceptance of the manuscript. The full R code and QIIME 2 script will be available on Zenodo upon acceptance of the manuscript.

### Author’s contributions

S.D. and B.M. conceived the study. A.G., C.D., S.D. and B.M. conceived the experiment. A.G. and C.D. conducted the experiment. S.D. and B.M. took part in the experiment. H.H. conducted histological data processing and analyses. A.G. and C.D. conducted molecular data processing. A.G., S.H., S.D. and B.M. analysed data. A.G., S.D. and B.M. wrote the manuscript. S.H. edited the manuscript. All authors contributed and agreed on the contents.

## Acknowledgements

Kandiah Santhirakumar managed fish maintenance, Claude Yéprémian advised on cyanobacterial cultivation and Arthus Escalas helped with statistical analyses.We thank the Amagen platform for providing medaka fish, ENVA for histological data processing, the PtSMB platform of MNHN for metabolomics.

## Additional Files

Supplementary figures and notes 1 and 2

Supplementary notes 1 and 2, figures S1 to S6, and Table S1.

Dataset S1

Sampling counts during the whole experiment.

Dataset S2

Monitoring of abiotic and biotic parameters during the experiment.

Dataset S3

Annotations of metabolites from the Microcystis aeruginosa strain PMC 728.11.

Dataset S4

16S rRNA gene sequencing and quality filtered read counts, and measures of alpha-diversity metrics (species richness, shannon, evenness).

Dataset S5

Annotations of the most correlated metabolites with bacteria from fish guts at day 28.

