## Supplementary note for "Disruption of fish gut microbiota composition and holobiont’s metabolome by cyanobacterial blooms"

### Supplementary Note 1

**Dataset pre-processing and bacterial community comparison within the entire sample set.** Overall, a total of 168 samples (4,235,404 reads) were obtained. After quality control, a total of 3,430,371 quality-filtered reads, corresponding to 3,198 Amplicon Sequence Variants (ASVs) was retained from a total of 164 samples including 5 replicates of *M. aeruginosa* cultures, 26 water samples, 15 bottom-growing biofilms, 1 fish food sample, 20 faeces and 97 whole gut samples (**see Dataset S3**). Eight ASVs representing at least at 1% of reads in the blank control sample were discarded, as well as 3 gut samples containing at least 3% of their reads coming from the blank control sample. After rarefying samples to 6,978 sequences, three samples were removed due to low read counts (food and two guts). The diversity of bacterial communities within compartments others than the gut has been examined to better grasp their dynamic (**Figure S1a-c**). Biofilms were the most diverse compartment (264 ASVs in average) and contained the most equitable bacterial communities (0.73 in average). *M. aeruginosa* cultures were much less diverse (41 ASVs in average), and water and guts much inequal (0.420 and 0.448 in average, respectively). As expected, the composition of bacterial communities was well differentiated with a quite similar composition between cultures, biofilms and water, unlike gut bacterial communities, and amid the faeces (**Figure S1d**).

### Supplementary Note 2

**Metabolic content of *Microcystis aeruginosa* cultures.** Biosynthetic gene clusters present in the PMC 728.11 *M. aeruginosa* strain genome (same strain as in this study) were retrieved from complete genome sequencing, as previously described (1), and were potentially involved in the biosynthesis of bioactive metabolites, such as various cyanopeptides, including mycosporine-like amino acids (MAAs), microcystins, cyanopeptolins, aeruginosins, aerucyclamides and bacteriocins. All were produced by the same PMC 728.11 strain used in this study, except aerucyclamides and bacteriocins although they were previously observed in PMC 728.11 cell extracts (2) (**Table S1 and see Dataset S3**). Among the annotated metabolites, microcystins, counting 11 different variants with MC-LR the most observed, amino acids and peptides were the most numerous metabolites produced by *M. aeruginosa* cultures.

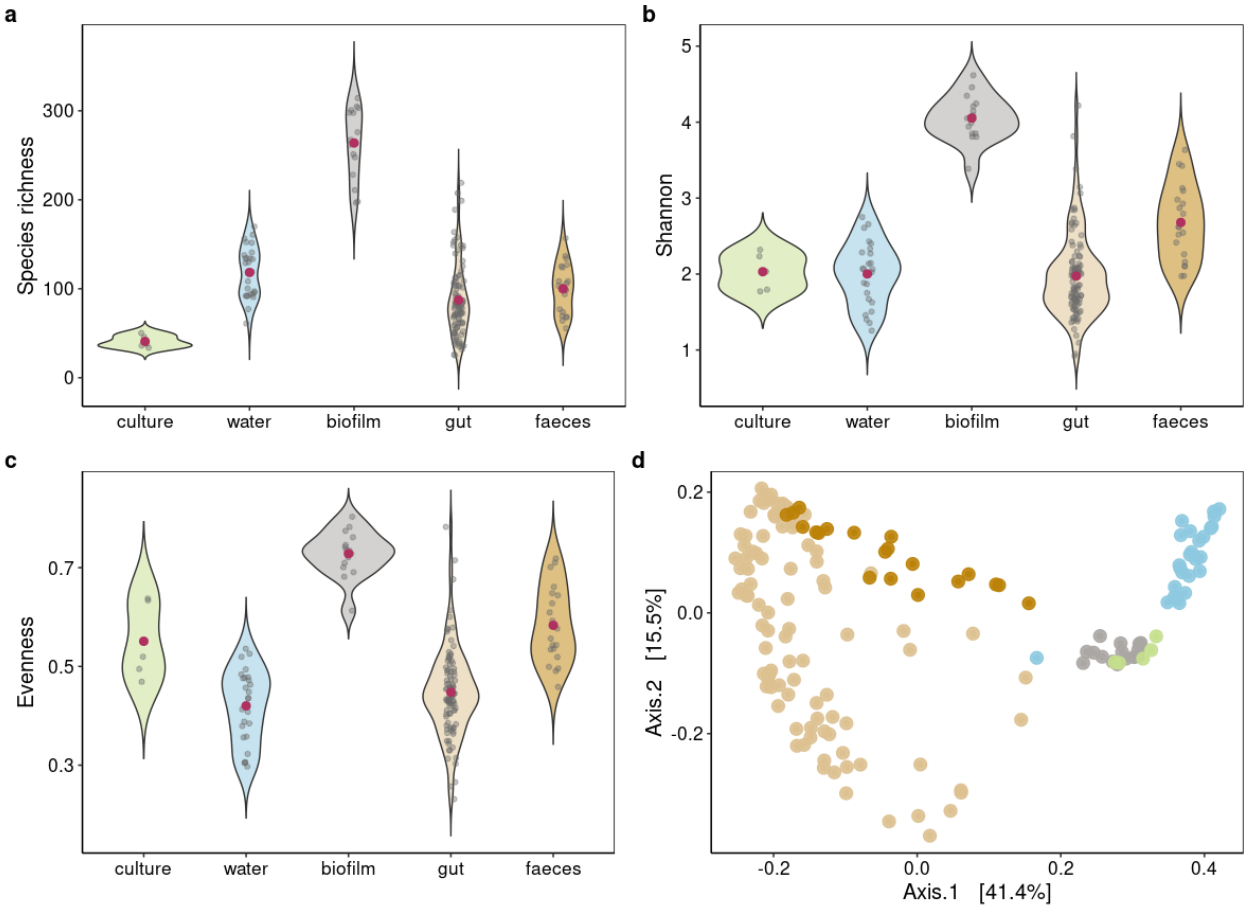


**Figure S1:** (a-c) Alpha-diversity metrics (species richness, Shannon, evenness) in the different sample types. Means are represented by a red dot. (d) Principal coordinates analysis (PCoA) representing bacterial communities from the different compartments using the weighted UniFrac distance.

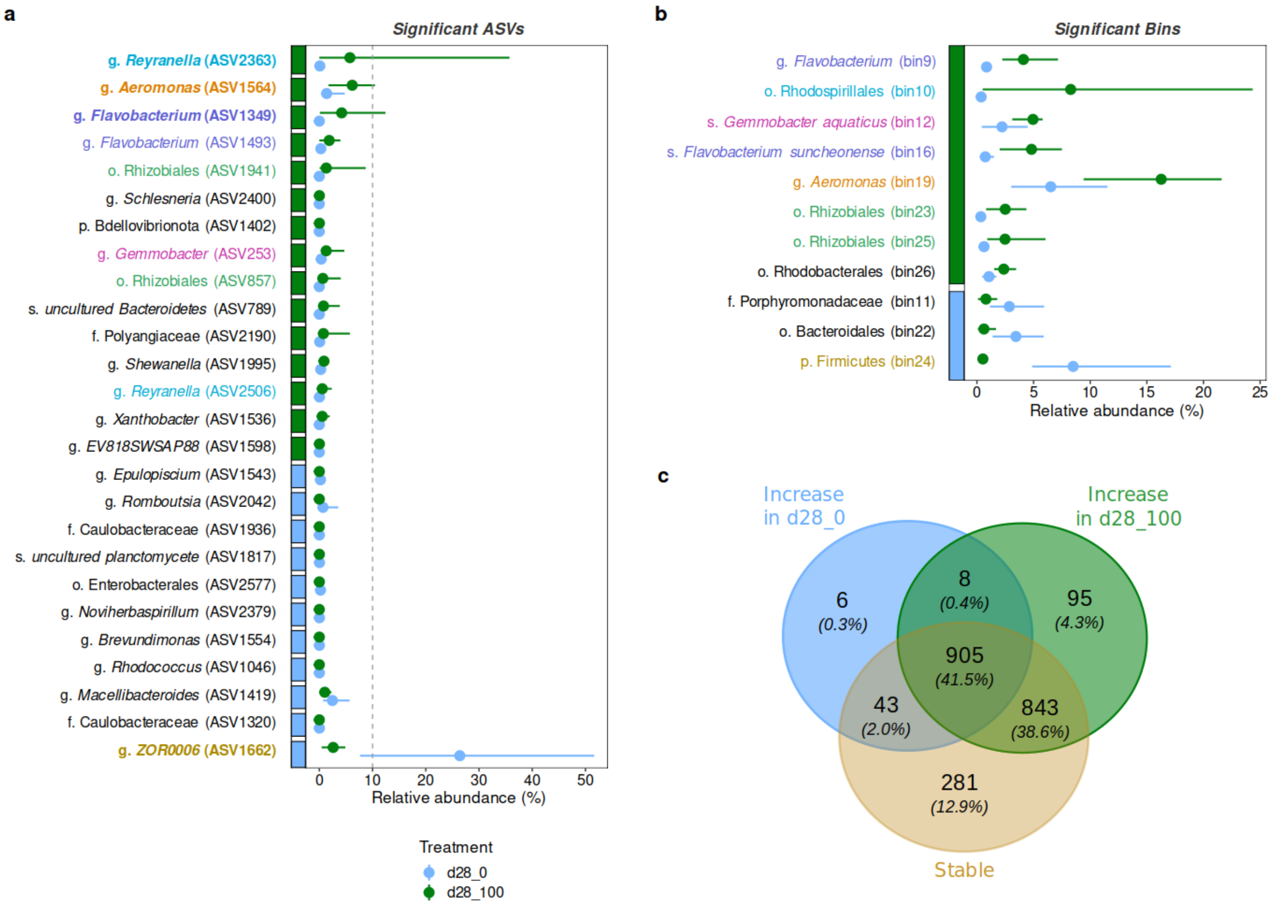


**Figure S2:** (a-b) Differentially abundant fish gut bacteria between treatments d28_0 (blue) and d28_100 (dark green) based on 16S rRNA reads (a) and shotgun metagenome sequencing (b). The absence of 16S rRNA sequences in metagenome bins does not allow to directly connect ASVs with bins, however the coloured taxon names represent bacteria with the same taxonomic affiliation. Lowest assigned taxonomic levels are displayed with the level specified by a single letter. (a) Discriminant ASVs based on the Linear discriminant analysis (LDA) effect size (LEfSe) with a LDA score above 3.5 and their variations in relative abundance in fish guts. The coloured boxes represent the treatment where each ASV is found the most abundant. The dots represent average relative abundance, the lines spread over the range of observed values. Only dominant ASVs (see text), were considered and further investigated. (b) Taxonomic affiliations (using the BAT method) of bins displaying significantly different relative abundances based on the Wilcoxon rank-sum test ($p < 0.05$) and their relative abundance across fish gut samples. The CAT method was used to affiliate bin10 (Rhodospirillales) at a lower taxonomic level, *i.e. Reyranella massiliensis*. The treatment where each bin is significantly more abundant is represented by the coloured box. (c) Overlap of KO counts among three groups: KO from bins significantly more abundant in d28_0 (blue) or in d28_100 (dark green) ($p < 0.05$, Wilcoxon rank-sum test), or KO from bins non-significantly different between the two treatments (beige) ($p > 0.05$).

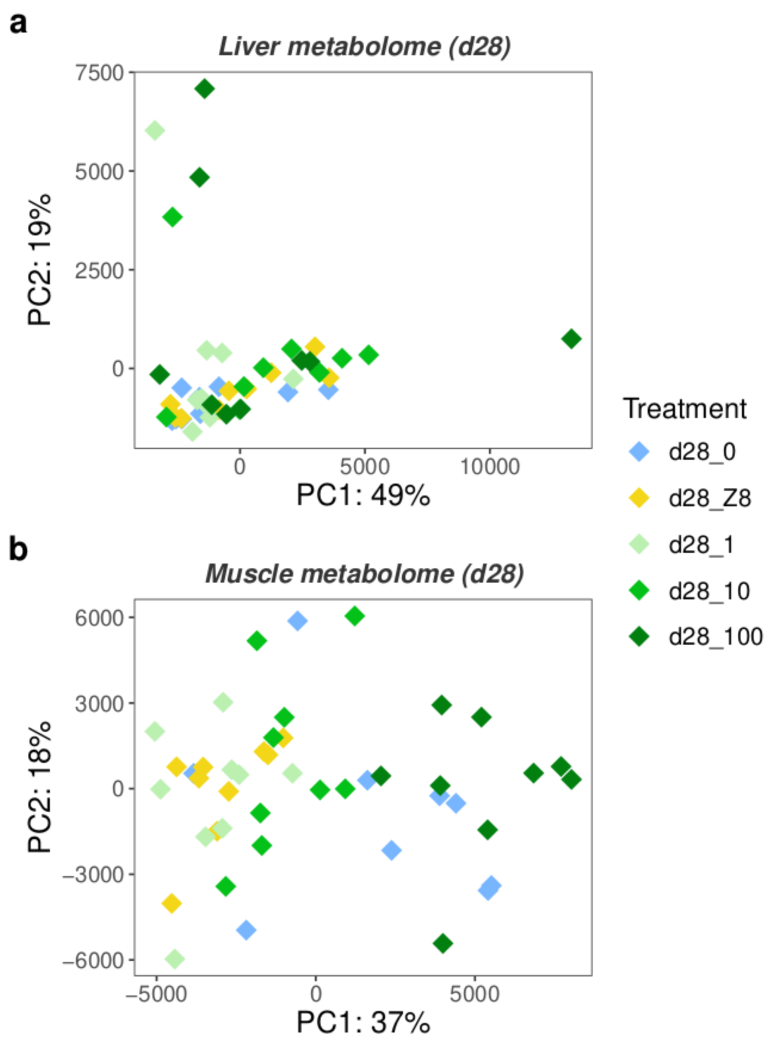


**Figure S3:** Principal component analyses (PCA) illustrating the metabolite composition in fish livers (a) and muscles (b) from the five different treatments during 28 days.

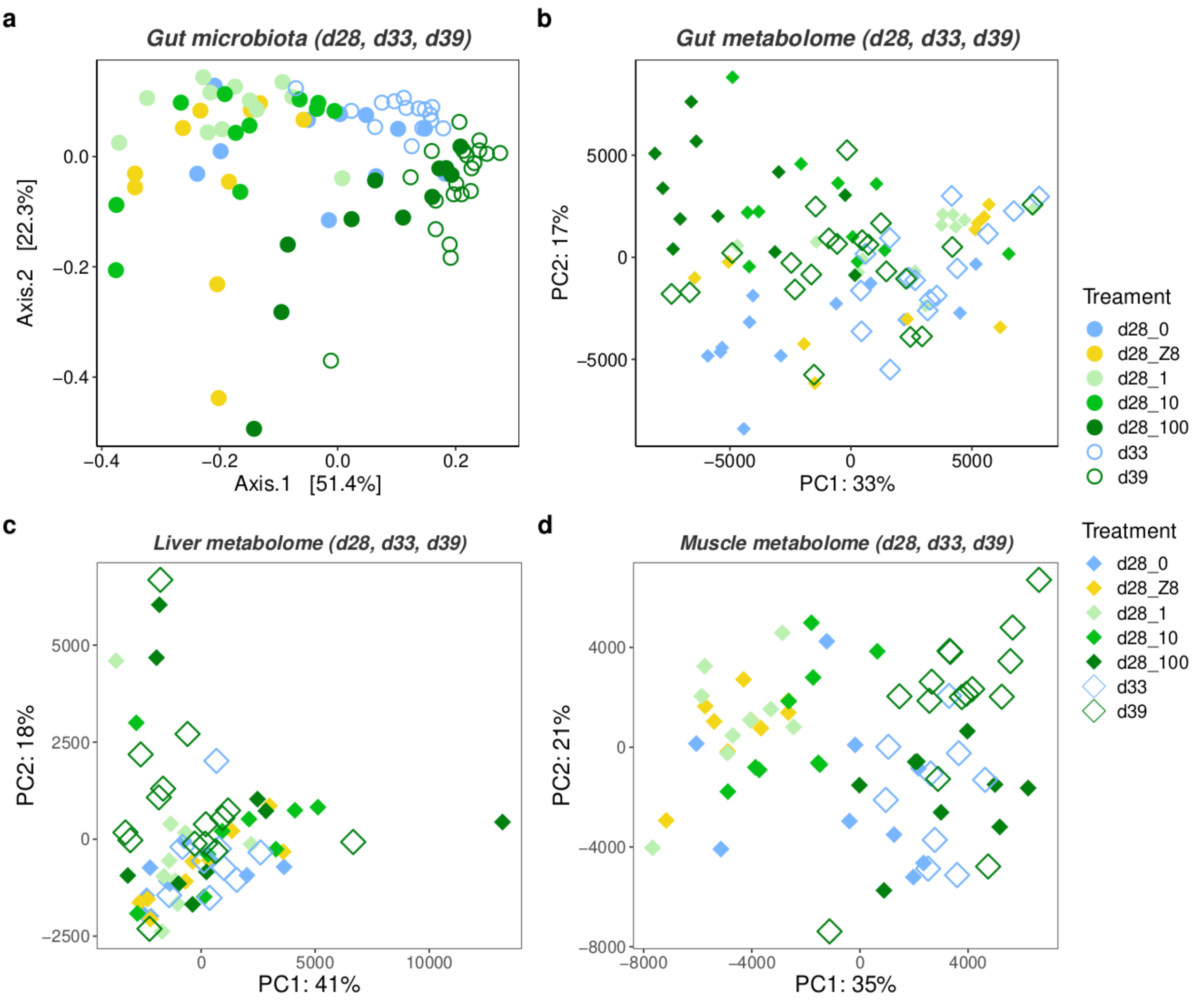


**Figure S4:** Comparisons of the composition of bacterial communities (a) and metabolite profiles (b) in fish guts, and the composition of metabolite profiles between the three different sampled organs, guts (b), livers (c) and muscles (d). The three organs were either exposed long-term during 28 (d28) days (full circles or diamonds) or short-term during 4 (d33) or 5 (d39) days (open circles or diamonds). (a) Principal coordinates analysis (PCoA) on weighted UniFrac distance illustrating bacterial composition in fish guts. (b-d) Principal component analyses (PCA) representing metabolite profiles in fish guts, livers and muscles.

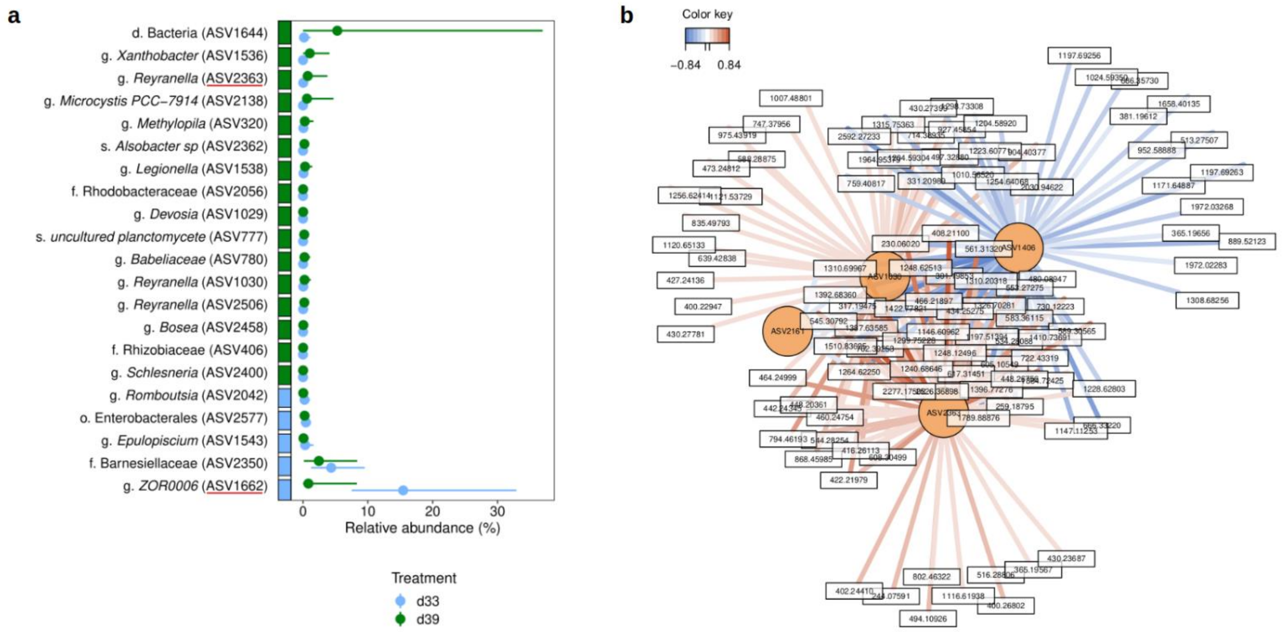


**Figure S5:** (a) Significant ASVs from the LEfSe analysis, differentially abundant between d33 gut samples (in water) and d39 gut samples (in 100 $\mu$g.L$^{-1}$ Chl*a*). The coloured boxes represent the treatment where ASVs are found more abundant. Dots and lines indicate mean and range of values, respectively. The displayed taxonomic affiliations correspond to the lowest assigned level, displayed using the first taxon letter. The two ASVs (ASV2363, ASV1662) underlined are also found differentially abundant between d28_0 and d28_100. (b) Relevance network analysis representing the most correlated ASVs and metabolites discriminating d33 and d39 gut samples. Only ASVs (in orange) and metabolites (in white) associated with Pearson correlation scores above ±0.714 are displayed. Coloured segments represent Pearson correlation values, either positive (red) or negative (blue).

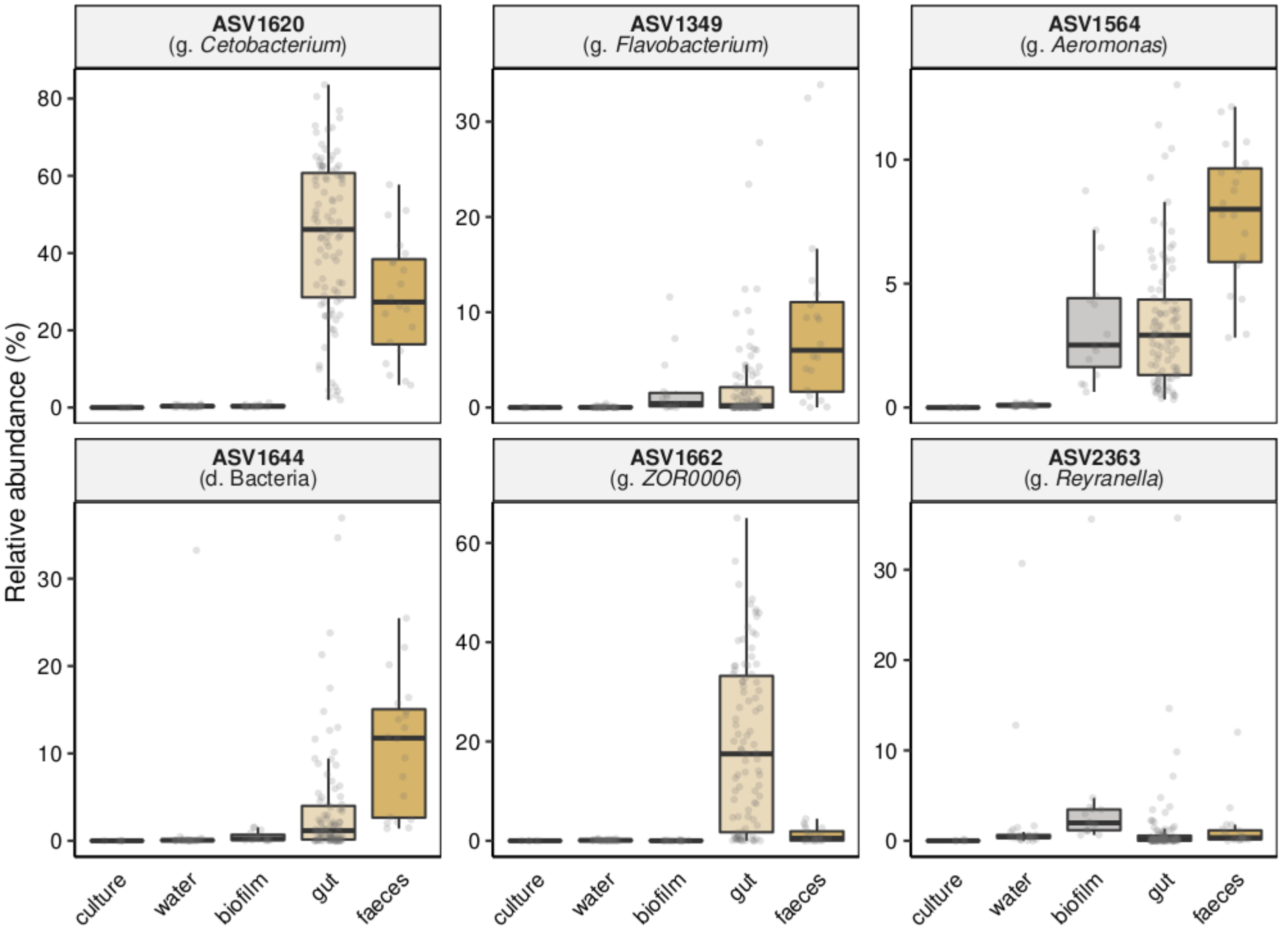


**Figure S6:** Relative abundance of dominant ASVs across compartments other than gut (culture, water, biofilm, gut and faeces). ASVs were searched for in the fish food sample discarded at the rarefaction step, but any ASVs were found (only 1% of reads for ASV1620).

| Metabolite annotation | Proportion (%) |
|---|---|
| Microcystin | 17.5 |
| AA/peptide | 15.9 |
| Phosphonic acid | 12.2 |
| Microcystbiopterin | 11.6 |
| Nucleic acid | 6.3 |
| Spumingin/aeruginosin | 6.3 |
| Lysophosphatidylcholine (LPC) | 4.8 |
| Benzenosulfonamide | 3.7 |
| Cyanopeptolin | 3.7 |
| Aminocyclitol glycoside | 3.2 |
| Microginin | 3.2 |
| Aeruginosin | 2.6 |
| Lipid | 2.6 |
| Mycosporine-like AA (MAA) | 1.6 |
| Cyanopeptolin-like AA | 1.1 |
| Lipopolysaccharide (LPS) | 1.1 |
| Macrolide | 1.1 |
| Benzenoid | 0.5 |
| Cyanopeptide | 0.5 |
| Phthalic anhydride | 0.5 |

**Table S1:** Composition of annotated metabolites of the PMC 728.11 *Microcystis aeruginosa* strain. Percentages represent the proportions of each annotated cluster on the total annotated clusters. Abbreviation: AA = Amino Acids.
